# Slow oscillations promote long range effective communication: the key for memory consolidation in a broken down network

**DOI:** 10.1101/2022.04.13.488133

**Authors:** Hamid Niknazar, Paola Malerba, Sara C. Mednick

## Abstract

One of the most prominent and robust findings in cognitive neuroscience is the strengthening of memories during non-rapid eye movement (NREM) sleep, with slow oscillations (SOs, < 1Hz) playing a critical role in systems-level consolidation. However, NREM is a sleep period generally showing a breakdown in connectivity and reduction of synaptic plasticity with increasing depth: a brain state seemingly unfavorable to memory consolidation. Here, we present a novel approach to address this apparent paradox that leverages an event-related causality measure to estimate directional information flow during NREM sleep in epochs with and without SOs. Our results confirm that NREM is generally a state of dampened neural communication, but reveals that SOs provide two windows of enhanced large-scale communication before and after the SO trough. These peaks in communication are significantly higher when SOs are coupled with sleep spindles, compared with uncoupled SOs. To probe the functional relevance of these SO-selective peaks of information flow, we tested the temporal and topographic conditions that predict overnight episodic memory improvement. Our results show that global, long-range communication during SOs promote sleep-dependent systems consolidation of episodic memories. A significant correlation between peaks of information flow and memory improvement lends a predictive validity to our measurements of effective connectivity. In other words, we were able to predict memory improvement on the basis of independent electrophysiological observations during sleep. This work introduces a non-invasive approach to understanding information processing during sleep, a behavioral stage whose function, until now, has been understood only after synaptic reorganization after waking from sleep. Our findings provide a mechanism for how systems-level brain communication can occur during an otherwise low connectivity sleep state, indicating that SOs are a gating mechanism for large-scale neural communication, which is the necessary substrate for systems consolidation and long-term memory formation.

## Introduction

Human brain oscillations measured by electroencephalography (EEG) reflect synchronized activity of thousands to millions of neurons, with spike timing modulated by the phase of ongoing oscillations (Legatt et al., 1980). Current models propose that oscillations, separately or in coordination with one another, serve to organize information processing and communication in neuronal cortical networks in a state-dependent manner (Buzsaki, 2004) and are thought to be markers of neural processing that occur in relation to specific behavior. A growing body of studies using neuronal recordings (R. Huber et al., 2008), cortical and local field potential (Mizrahi-Kliger et al., 2018), EEG signals, cognitive and behavioral methods (Naji et al., 2019) have demonstrated the important and causal role of non-rapid eye movement (NREM) slow oscillations (SOs, <1Hz) in brain function.

Slow oscillations are generally understood as large travelling waves that coordinate activity globally across cortical regions and in cortical-subcortical interactions (Malerba et al., 2019; Massimini, 2004) and have been shown to support memory formation (Marshall, 2004; Marshall et al., 2006). According to the framework of systems consolidation, long-term memories are initially bound by a fast-learning system in the hippocampus (i.e. encoding), followed by stabilization of these memory traces in cortical stores during NREM (i.e., consolidation) (Diekelmann & Born, 2010a). One potential way NREM may accomplish this process is by synchronizing neuronal activity along cortico-thalamo-hippocampal circuits via global SOs that coordinate nested spindles (sigma power; 12-15Hz) and sharp wave ripples (SPW-R) (Born, 2010; Ngo et al., 2020; Niethard et al., 2018; Rasch & Born, 2013). Indeed, triple phase coupling of these rhythms is associated with memory improvement and causal interventions that block or enhance SO activity alter memory performance, suggesting a critical role of SOs in this process (Ramadan et al., 2009).

This prominent hypothesis of sleep-dependent memory, which posits increased activation of global communication networks, stands in opposition to the known properties of NREM sleep, which is characterized by reduced plasticity and a loss of network communication. Studies have shown that long-term potentiation, a form of synaptic plasticity that is the leading physiological model for the initial encoding and subsequent stabilization of memory, is actively suppressed during NREM sleep (Jones Leonard et al., 1987), compared with REM sleep or an anesthetized state (Bliss & Lømo, 1973). At the systems level, using a combination of transcranial magnetic stimulation and high-density EEG, Massimini and colleagues demonstrated a breakdown of transcallosal and long-range neural communication as the brain transitions from wake to the deeper stages of NREM sleep (Massimini et al., 2005), suggesting that network communication during NREM sleep is highly local. Thus, the fundamental properties of NREM sleep (reduced synaptic plasticity, local, disconnected network) and the hypothesized mechanism of SO-driven systems consolidation (coordinated, global network activation) appear contradictory and cannot explain how NREM supports memory consolidation. From a theoretical perspective, the kind of memory consolidation we consider here rests upon long-term potentiation or depression; namely, activity-dependent plasticity. The requisite synaptic activity depends upon directed (effective) connectivity; i.e., ongoing communication in neuronal networks. Current formulations of this plasticity speak to long-term depression and synaptic regression. This view emerges from both empirical studies (Gilestro et al., 2009; Reto Huber et al., 2004; Tononi & Cirelli, 2006) and theoretical considerations based upon optimizing generative models (Hinton et al., 1995; Hobson & Friston, 2012). In these accounts, emphasis is placed upon removing redundant or exuberant synaptic connections during sleep to ensure generalization; i.e., retaining the associations that matter.

The current study addresses the apparent paradox of NREM sleep supporting plastic activity-dependent changes in a globally coordinated network by investigating directional information flow during NREM sleep epochs with and without SOs. Prior studies in humans have typically assessed the functional role of sleep in memory at a distance, whereby memories are encoded, followed by a subsequent sleep period, after which memories are retrieved. This strategy does not allow for understanding of how the ongoing processing of sleep directly affects the consolidation of memories. To take a first step in amending this gap, we have adopted an event-related approach to measure causal information flow across the brain during NREM sleep, specifically comparing different phases of the SO with non-SO windows. In addition, studies typically analyze SOs using functional connectivity to measure temporal similarity or correlations between different EEG channels (Fox & Raichle, 2007), which is limited to correlational measures, and cannot identify directional causal communication. In contrast, we leverage effective connectivity, which tests the influence that one neural system exerts over another either directly or indirectly (Friston, 2011) estimated using Granger causality. According to this approach, a causal relation is detected if past values of a primary signal (source) help predict a second signal (sink) beyond the information contained in its past alone (Granger, 1969). Thus, the current study non-invasively probes how the brain organizes network activity during sleep to support memory, by measuring event-related causal information flow across EEG channels during the SO.

### Causal information outflow from sources during SOs

We used generalized partial directed coherence (GPDC) (Baccala et al., 2007) as an estimator of causal information flow between different EEG channels. GPDC was defined as an extension of partial directed coherence (PDC) (Baccalá & Sameshima, 2001) to make it more robust against the amplitude scale of time series (see ‘Effective connectivity estimation’ section for details). Four EEG channels (Fz, Cz, Pz, POz) were considered as the source of information flow and 12 channels (three channels in each region of frontal, central, parietal and occipital), were considered as the sink of information flow. An automatic system detected SOs in the source channels and two quantifiers were computed in the window 1sec before to 1sec after the SOs trough (Figure 1.A). We defined the first quantifier as the outflow from a source (*CH*_*outflow*_, ‘channel outflow’) and the second quantifier as flow from a source channel to a sink region (*CH*→*R*). Both quantifiers were estimated with respect to the SO phase and computed as an average for each subject (see Methods for definitions and details). Figure 1.B shows the average and standard deviation of *CH*_*outflow*_ for each source across SO channels and based on SO phase. As SO phase changes, outflow peaks at 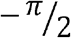 (a ‘pre-peak’), drops to a local minimum at 0 (corresponding to the SO trough) and peaks again at (a ‘post-peak’). As a comparison, we also estimated outflow from sources during non-SO windows. We selected 500ms-long time windows for all SOs in a randomized manner, values were chosen among the four sources channel and in either sleep stage (S2 and SWS) with the only condition that the time epochs had to be more than 10sec away from any SO trough. Figure 1.C displays the outflow values for each source in pre- and post-peaks, the SO trough, and non-SO windows. We tested outflow differences in SO and non-SO windows with ANOVA and post-hoc analysis. The results showed that outflow () in the pre- and post-peaks was significantly larger than outflow in non-SO windows (p-value<0.001), but outflows at SO troughs and the non-SO windows were not different (p-value>0.05) (Figure 1.C). Also, compared to non-SO windows, there was no significant difference between outflow in half a cycle after SO trough (phase =) and significant or marginally significant (0.05<p-value<0.1) difference between outflow in - phase, which can be caused by the effect of a prior SO on the successive SO in a sequence of SOs.

**Figure 1.**
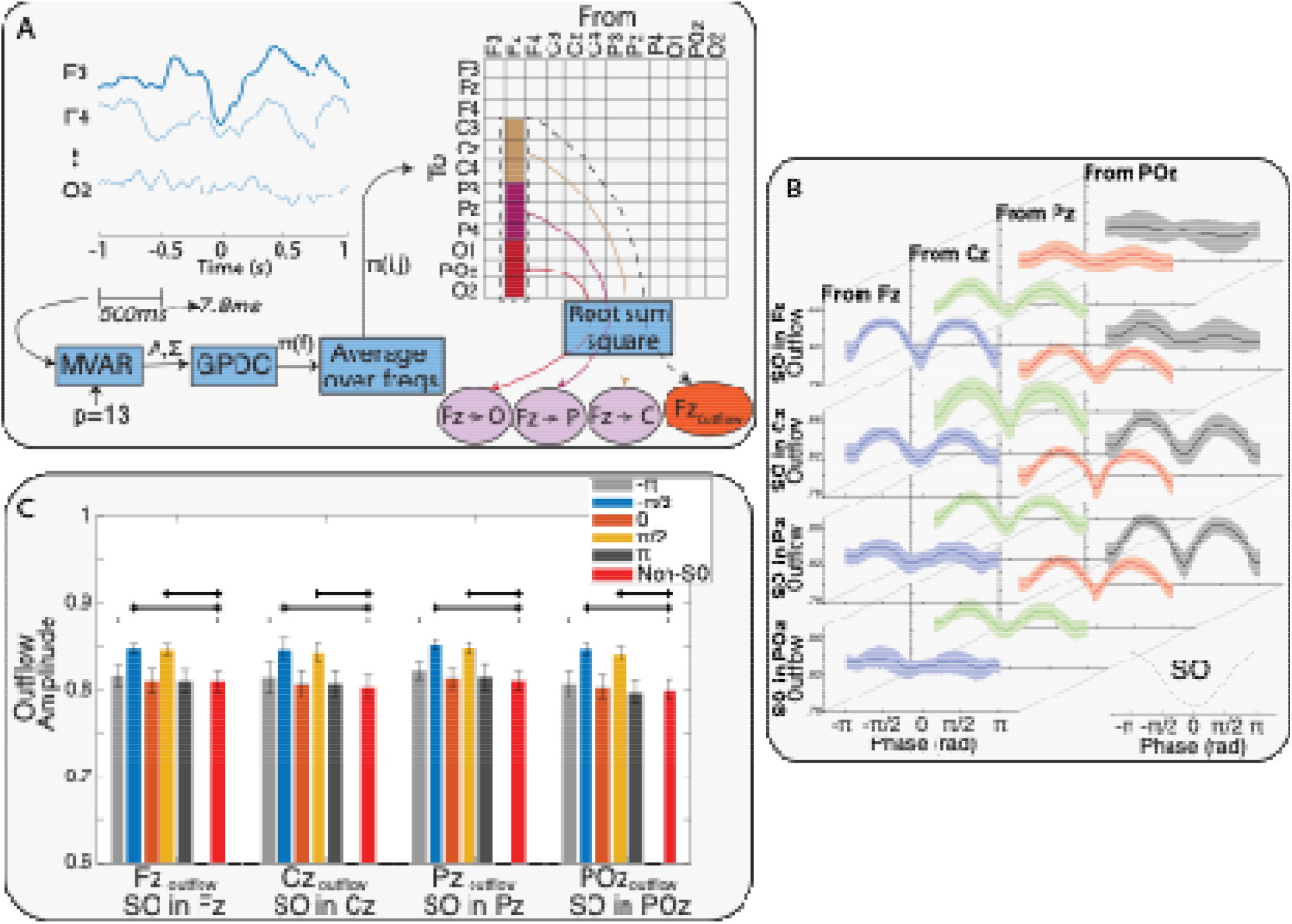
A. A representation of the process that estimates causal information flow between EEG channels. In summary, after detecting SOs, a multivariate autoregressive model was fit on each 500ms window of 12 EEG channels from 1sec before to 1sec after each SO trough, with stride of 7.8ms. By using the GPDC method and averaging over all frequencies, the effective connectivity matrix () and the two defined quantifiers were calculated. After resampling the quantifiers time series based on the SO phase, the phase series were averaged over all SOs of each subject to obtain the phase series of 1) the outflow from the sources (e.g., Fz_Outflow_) and 2) the flow from sources to sinks (e.g., Fz→Cz) for each subject. B. Variation of outflow from all sources conditioned on the SO channel. Each row shows the outflow variation across the subjects over phase of SOs when the SO was detected in the represented channel. In Each row, the columns show outflow variation in each of the sources (mean ± std). The phase series and their variation across the sources and SO channel show peaks before and after the SO trough and a possible relation between source and SO channel. The results in this figure suggest that sources closer to SO channels had greater height in outflow peaks. C. Comparing the outflows at five different phases of the SOs 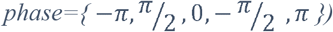 to outflow in non-SO windows. ANOVA and post-hoc tests found that peaks in phase of 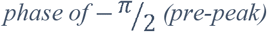 (pre-peak) and 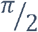 (post-peak) had significantly larger outflow, while there were no significant differences between outflow in non-SO windows and SO trough. Thick black lines indicate a strong significant difference (p-value<.005), thin lines indicate significant difference (p-value<0.05) and dash lines indicate marginal difference (0.05<p-value<0.1) between SO and non-SO windows.

To investigate if high amplitudes during SOs were driving these peaks in information flow, we tested if there were correlation between peaks of information outflow and amplitude of SOs in their relevant time point. The results showed for each subject there were no significant relations between SO trough amplitudes in each channel and the outflow peaks from the relevant channels (Figure S.2). Also, there were no significant linear relations between SO amplitude averages in each channel and average of the outflow peaks from the relevant channel (Figure S.3).

We then asked if there was a structure underlying how high the peaks of information flow will be based on the various elements that allowed their estimate (locations of sources and sinks, etc.). To estimate the relative contribution to peak height of all these elements in an un-biased and comprehensive model, we fit a linear mixed-effects (LME) model to examine the effect of sources, SO channels, and distances between sources and SO channels 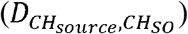 and peak phase (i.e., whether the peak preceded or followed the SO trough) on the height of the peaks in information outflow. We assigned discrete values for the distances 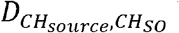 between source (Fz, Cz, Pz and POz) and SO channel as zero to three in the linear model, adding one for each ‘step’ in the frontal-occipital axis. We used a simple model to calculate the distances as we considered distance between each two neighbor regions equal to 1. (This meant that if *CH*_*source*_ was Fz and was Cz, difference would be 1, and if *CH*_*suorce*_ was Cz and *CH*_*so*_ was POz, the distance would be 2). Table S.1 presents the coefficients and related p-value identified by the LME model. Also, we added SO amplitude to the model to test the effect of this predictor on the height of outflow peaks. Adding SO amplitude did not improve the model, implying that there was no relation between SO amplitude and peaks of information outflow (P-value>0.05).

Based on LME modeling outcome, there were significant linear effects of the SO channel (p-value<0.05), distance between the SO channel and the source 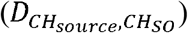, and phase (pre-/post-peak) on the height of outflow peaks (p-values<0.05), 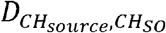 with, having the largest linear effects (higher coefficients). The negative coefficient of 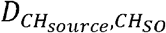 demonstrated that information outflow increased with proximity of the sources to the SO channel. For example, for an SO detected at the Fz channel, the lowest peak of causal information outflow would be found in the POz channel, whereas moving closer to the Fz channel, the peak of the outflow would increase, with highest outflow at Fz itself. The positive coefficient of the SO channel indicates that the position of the SO (anterior vs posterior) modulated the height of the outflow, where SOs in the anterior channels may produce lower outflow in all sources compared with SOs in more posterior regions. Additionally, the negative coefficient of peak phase (i.e., being a pre- or a post-peak) indicates that pre-peaks are expected to have greater outflow than post-peaks.

### Causal information flow from sources to sinks during SOs

In the previous section we showed that the SO-related information outflow from a source was significantly affected by the distance between the SO channel and the source, where sources closest to the SO had largest peaks. To analyze how information flow depended on sender and receiver, we next focused on source – sink pairs. Causal information flow from each source to each sink (*CH* → *R* quantifier, *FZ*→ *P* where would have Fz as source and parietal as sink) were computed over all SOs. For each SO channel, we averaged all *CH* → *R* phase series in each source–sink pair within each subject. Figure 2.A presents samples of *CH* → *R* phase series variation from two of the sources (Fz and POz) to three sinks separately (central, parietal and occipital with Fz source and frontal, central and parietal with POz source), all cases with SO channel in Fz (see Figure S.5 to S.8 for comprehensive results).

**Figure 2.**
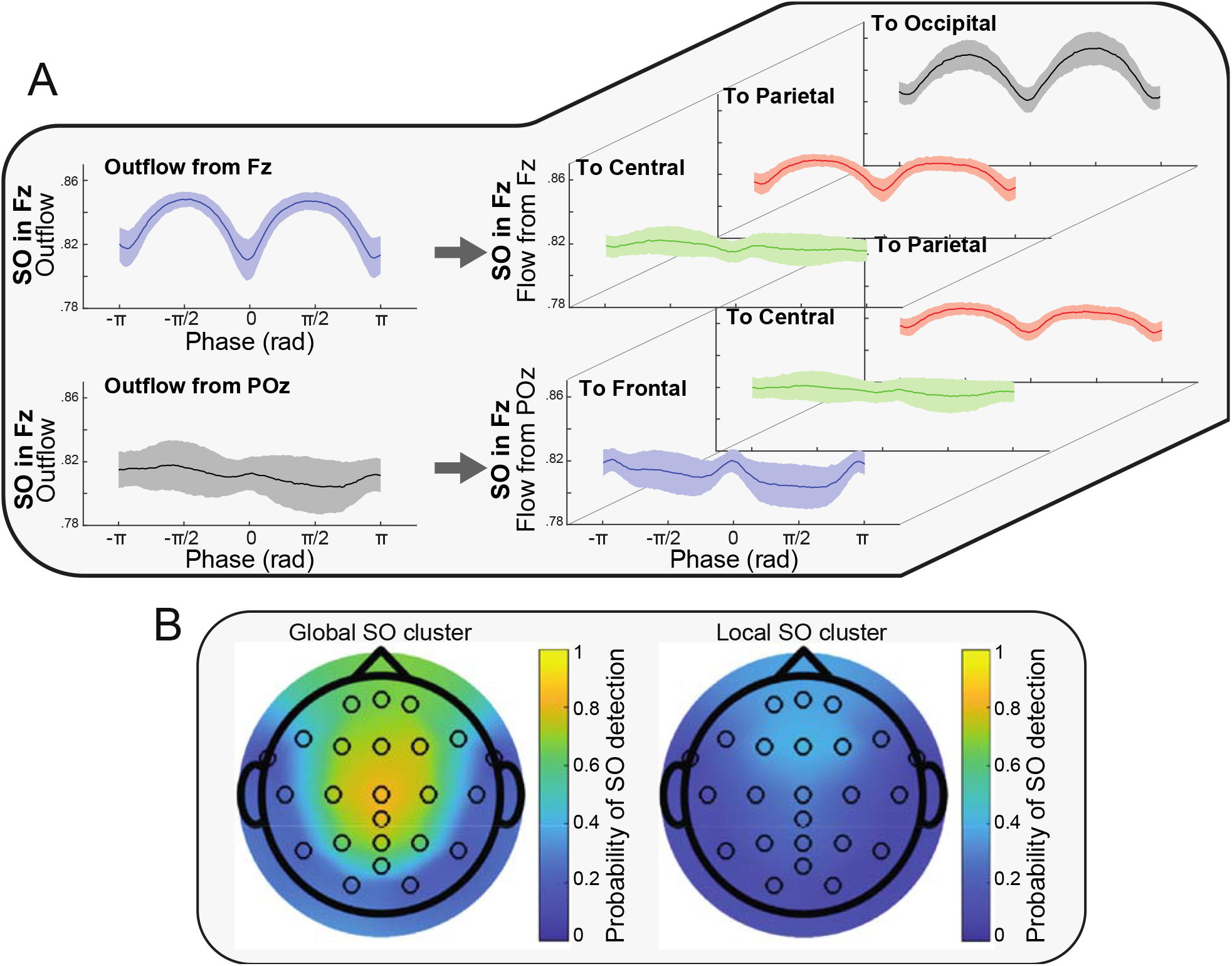
A. Examples of information flow from source to sink (CH → R quantifier). The left plots show the total outflow from the specified source and with presence of SO in the specified channel (Samples from Figure 1.B). The right plots represent the portions of the flow from the source to the different sinks. For example, the top-left plot shows the outflow from the Fz as the source when there were SOs in Fz channel and the three top-right plots represent portions of that flow to each of the sinks. By considering the topology of the SO channel, source and sinks the results suggest a relation between distance of sink to SO channel and the peaks of information flows. B. Two clusters of SOs found by the clustering method (Cluster 1: Global SOs, Cluster 2: Local SOs). The colors represent the density of the SOs in each channel.

Observing the *CH*→ *R* quantifier values in Figure 2.A (and Figure S.5 to S.8), we noticed that there was a relation between distance of the sink to SO channel 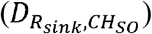 and height of the flow peaks. This suggested that the causal information flow (*CH* → *R*) could be modulated by which source–sink pair was considered, the distance of source–sink pairs to the SO channel, and the distance between the source and the sink in each pair. To investigate these potential effects, an LME model was fitted to the values in the peaks of the flow (pre- and post-SO trough), with the following linear predictors: SO channel, source–sink pairs, distance of source– sink pairs to the SO channel 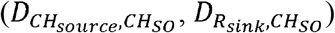 and distance between source–sink pairs 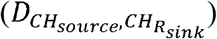.

The outcome of our LME model (Table S.2) demonstrated a significant linear effect of 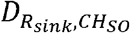 and 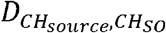 on the height of the flow peaks. Also, there was significant effect of peak phase (pre- or post-trough) where peak height in the pre-trough phase was larger than in the post-trough phase (coefficient<0). In contrast, we found no statistically significant linear effects of SO channel, source, or distance between source–sink pairs on the causal information flow peak height. The coefficient values showed different magnitude and direction of the effect in each predictor. The positive coefficient of distance between sink and SO channel and negative coefficient of distance between source and SO channel revealed that sources closer to the SO channel (less distance) and sinks farther from the SO channel (larger distance) had higher peak of information flow in comparison to other compositions. For example, for an SO in the *Fz* channel, sources closest to the SO channel and sinks farthest from the SO channel (e.g., FZ → 0) have the highest information flow, suggesting that brain communication during the SO is highest at locations far from the SO, and distance between SO channel to the source–sink pair impacts causal flow.

Our results support the following:

- Sources closer to the SO channel send the greatest magnitude of information (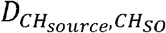, coefficient for flow peaks <0).
- Sinks farther from the SO channel receive the greatest magnitude of information (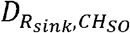, coefficient for flow peaks > 0).
- The distance between the sink and the SO channel shows strongest effect on information flow compared with all other predictors (absolute value of 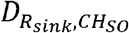 coefficient for flow peaks >> absolute value of other predictor coefficients).

### Effect of SO clusters on causal information flow

So far, we have shown that the impact SOs on causal information flow depends on the distance between the source to the SO channel when considering outflow, and depends on the distance of the sink to the SO channel when looking at specific flow between source-sink pairs. Also, channels closer to the SO channel can send more causal information flow, whereas channels farther from the SO channel can receive more causal information flow. In light of these properties, we hypothesized that the primary benefit of SO’s effect on brain information processing is to permit causal communication between remote brain regions. If this was true, it follows logically that global SOs (SOs that propagate across a large portion of the scalp) should mediate more information flow compared to other (more localized) types of SOs. Studies investigating the spatial and temporal co-occurrences of SOs report that the majority of SOs are traveling waves, with their pattern of origin and propagation providing an outline of cortical connectivity (Massimini, 2004). Using a cluster-based analysis of SO co-occurrences across the EEG channel manifold, prior work from our group identified three spatio-temporal categories: frontal, local and global (Malerba et al., 2019), whereby global SOs occur in most electrodes, local SOs occur in few electrodes without location specificity, and frontal SOs are confined to the frontal region. Along with having greater footprint and larger amplitude, global SOs also provided greater hierarchical nesting of thalamocortical sleep spindles, suggesting functional differences between SO categories, with global SOs more poised to activate systems consolidation (Malerba et al., 2019).

We tested the hypothesis that global SOs should have greater causal flow than non-global (local) SOs by comparing the effective connectivity in each type. We first clustered global and non-global SOs using a method previously introduced by our team (Malerba et al., 2019). Figure 2.B shows the occurrence rate of SO in each of the channels and in each of the two clusters on the scalp surface after the clustering process. Based on results from (Malerba et al., 2019) and our presented results in Figure 2.B, the two detected clusters were interpreted as global and local SOs clusters. To test the effect of cluster on amplitude of SOs, first we tested if there was a significant difference between amplitude of SOs in the global and local clusters. The results (Figure S.4) showed significant differences in the clusters in each channel (p-value<0.05). Then, we tested if in either cluster there was a significant relation between amplitude of SOs and height of information outflow peaks. Our tests showed no significant relation between SO amplitudes and peaks of information outflow. Our tests showed no significant relation between amplitude of SOs and peaks of information outflow. To test if there was an effect of cluster identity (i.e., being a global or local SO) on the height of information flow peaks, we added cluster identity (global as 1 and local as 2) as a fixed effect along with other used predictors in an LME model of information flow peak height.

The results of LME models for peaks of information flow from the sources to the sinks (Table S.3) showed the same effects of the previous predictors combined with a significant linear effect of clusters on the height of the flow peaks (p-values < 0.05). Negative coefficients for the cluster effect showed that peak heights were higher in global SOs (cluster = 1) than local SOs (cluster = 2). We interpret this as confirming our hypothesis that global SOs mediate larger information flow compared to other SO types.

As a complementary analysis, we tested effect of SO-spindle coupling on causal information flow by adding coupling as a predictor to the LME model. First, we detected all spindles which had overlap with the time window of ±1sec of SOs trough. We used the same approach as(Zhang et al., 2020) for spindle detection. Then, we added SO-spindle coupling as a fixed effect to the LME model by considering the value of non-coupled as 1 and coupled as 2. The result of modeling peaks of causal information flow by adding SO-spindle coupling showed a significant (p-value<0.05) positive effect of coupling (Table S.4), with the positive coefficient for the coupling effect indicating that peak heights were higher when SOs were coupled with spindles.

### Long-term memory improvement and causal information flow during SOs

Next, we investigated if SO properties that we identified as crucial for information flow has functional consequences on the formation of long-term memories. Based on the importance of down-to-up state transitions (Cousins et al., 2014; Göldi et al., 2019; Ngo et al., 2013; Rihm et al., 2014; Schreiner et al., 2018), matched to the post-peak in our study, we examined the relation between word-pair association (WPA) improvement and causal information flow calculated during the post-peak (peak of information flow at 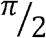 after the SOs trough). WPA improvement was measured as the ratio of performance after the sleep night (post-sleep) to the performance before the sleep night (pre-sleep). For each subject, we assigned the post-peak outflows to one of the four distances between the source and the SO channel 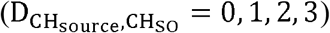 and then averaged within each group. We then evaluated linear relations between the outflow in each group and WPA improvement using linear regression models (Figure 3).

**Figure 3.**
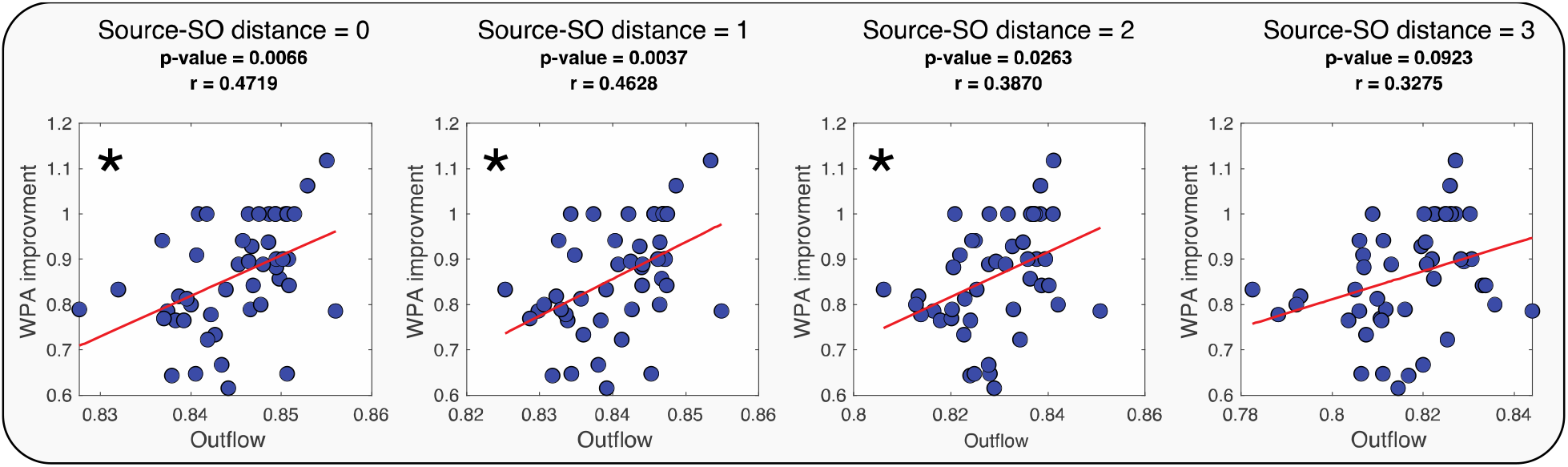
Regression test for outflow of source and WPA improvement in four different conditions of distance between SO channel and outflow source 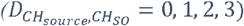. In each, we show the p-value, the linear coefficient is reported as r and the significant linear relationships are marked with asterisks (with p-values adjusted to Bonferroni correction).

Results in Figure 3 show that memory improvement is associated with the distance between SO and sources, with a significant positive linear relation between WPA improvement and outflow when the source is close to the SO channel 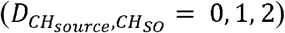, but not when the source is far from SO channel 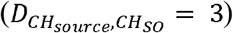. 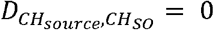 had the greatest r-value in the regression test and the r-value got smaller as distance between the channels increased 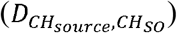: this can be interpreted as the outflow from sources close to SO was a better predictor of WPA improvement.

Next, we examined how the distance between source and sink channels, and the relative distance between the source and sink to the SO channel, impacted WPA improvement (Figure 4.A). First, we defined three possible distances between sink and source 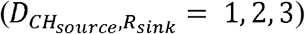 and averaged flows from sources to sinks which had the same distance. Figure 4.B displays the linear relation tests of WPA improvement and averaged flows in each specified distance between sink and source.

**Figure 4.**
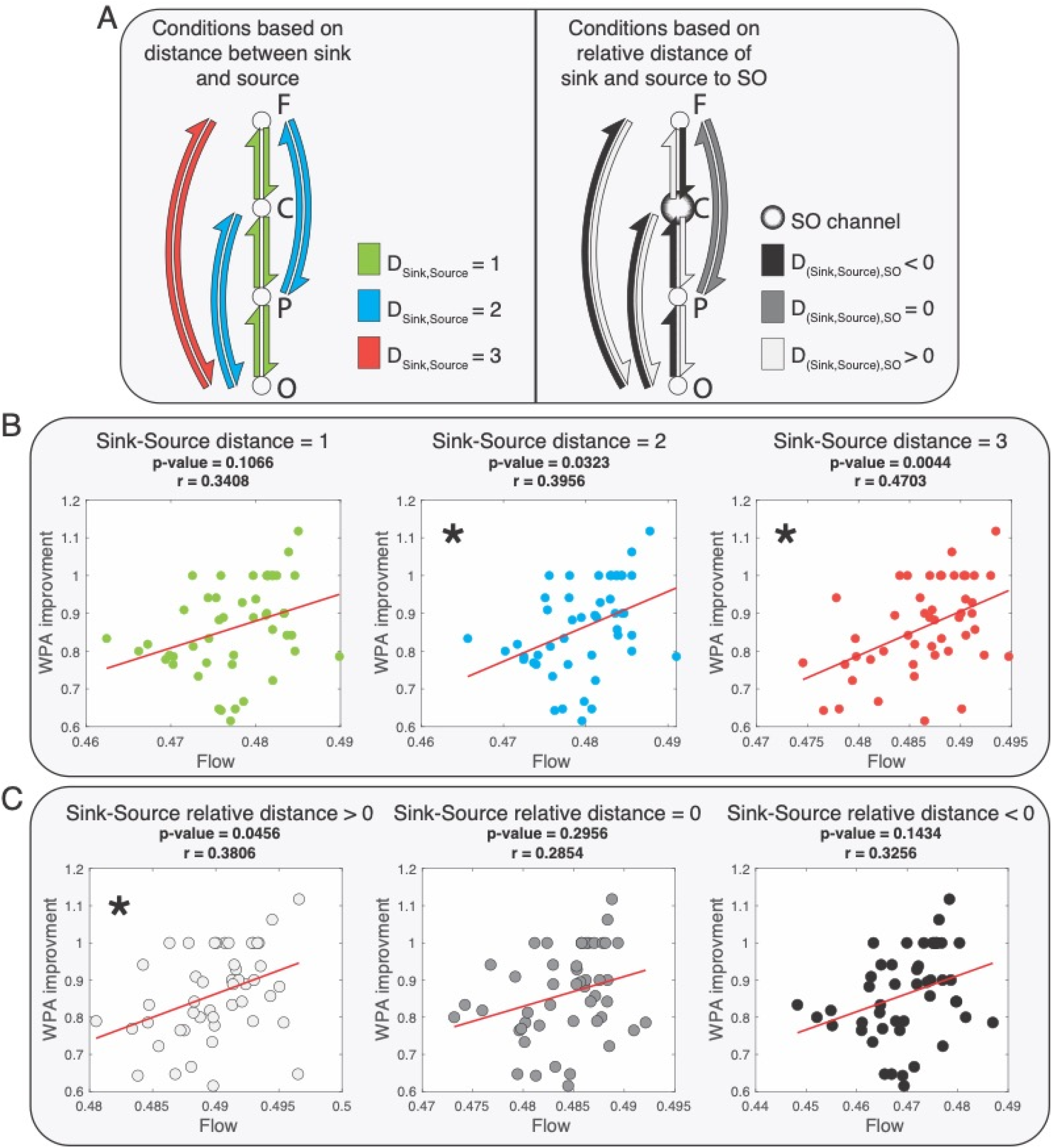
Linear relationships between flow and WPA improvement based on different conditions of distance between sink and source, and relative distance of sink and source to SO channel. The asterisks mark significant linear relationships (with p-values adjusted to Bonferroni correction). A. A representation of the three conditions of distance between sink and source (left graph) and an example of three possible conditions of relative distance of source and sink to SO channel (right graph, for SO channel at Cz). In the left graph each color represents pairs of sink and source of information flow with the same distances. The right graph shows an example of the conditions of the relative distance when the SO channel is Cz. The relative distance is greater than 0 when source is closer to SO channel than sink to SO channel and smaller than zero when sink is closer to SO channel than source to SO channel. B Results of correlation and regression test of relation between the flow and WPA improvement for three conditions of distance between source and sink 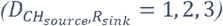. C. Results of correlation and regression test of relation between the flow and WPA improvement for three conditions of relative distance between source and sink to SO channel.

Based on the results presented in Figure 4.B, larger distances between sinks and sources 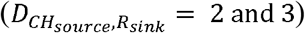 were linked to significant positive linear relation between information flow and WPA improvement. The r-value increased with greater distance between sink and source, suggesting a stronger linear relation between information flow and WPA improvement for long-range communications. The use of the correlation coefficient as a summary statistic - encoding the relationship between effective connectivity and memory improvement - scores the reliability of this association, as opposed to its effect size. In other words, the effect of distance between sources and sinks is on the correlation between connectivity and memory. This could reflect an increase in the effect of connectivity on memory - or a decrease in noise. This contrasts with an analysis testing for the interaction between peak connectivity and distance measures in predicting memory improvement.

Next, we examined the association of WPA improvement and flow in different relative distances from the sink and source to SO channel. We defined the relative distances as the difference of distance between sink and SO channel and distance between source and SO channel 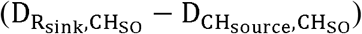. Hence, relative distances greater than zero indicated that the source was closer than the sink to the SO channel. For example, if Cz was the SO channel, the relative distance of Fz source and occipital sink would be greater than zero, as 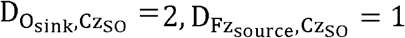. Figure 4.C shows a significant linear relation between WPA improvement and causal information flow only when the relative distance is greater than zero. Together, these results are consistent with our proposed hypothesis that long-range communication mediated by SOs plays an important role in memory consolidation. Also, to test if there is a relation between amplitude of SOs and WPA improvement, we correlated SO amplitude with and without considering the SO’s clusters. The results (Figure S.9) showed no significant relation between amplitude of SOs and WPA improvement (p-values < 0.05).

A final test of this hypothesis investigated whether global, but not local, SOs mediated the relation between causal information flow in SOs and WPA improvement. We first measured outflow in the post-peak SO phase within global and local SO clusters and calculated significance levels of regressions between outflows and WPA improvement, shown in Figure 5.C (refer to supplementary for the detailed results, Figure S.10 and S.11). We found a greater number of channels with a significant linear relation between their outflow and WPA improvement in the global cluster compared with the local cluster (4 combinations of source and SO channel in the local cluster and 13 combinations of source and SO channel in the global cluster, Figure 5.C).

**Figure 5.**
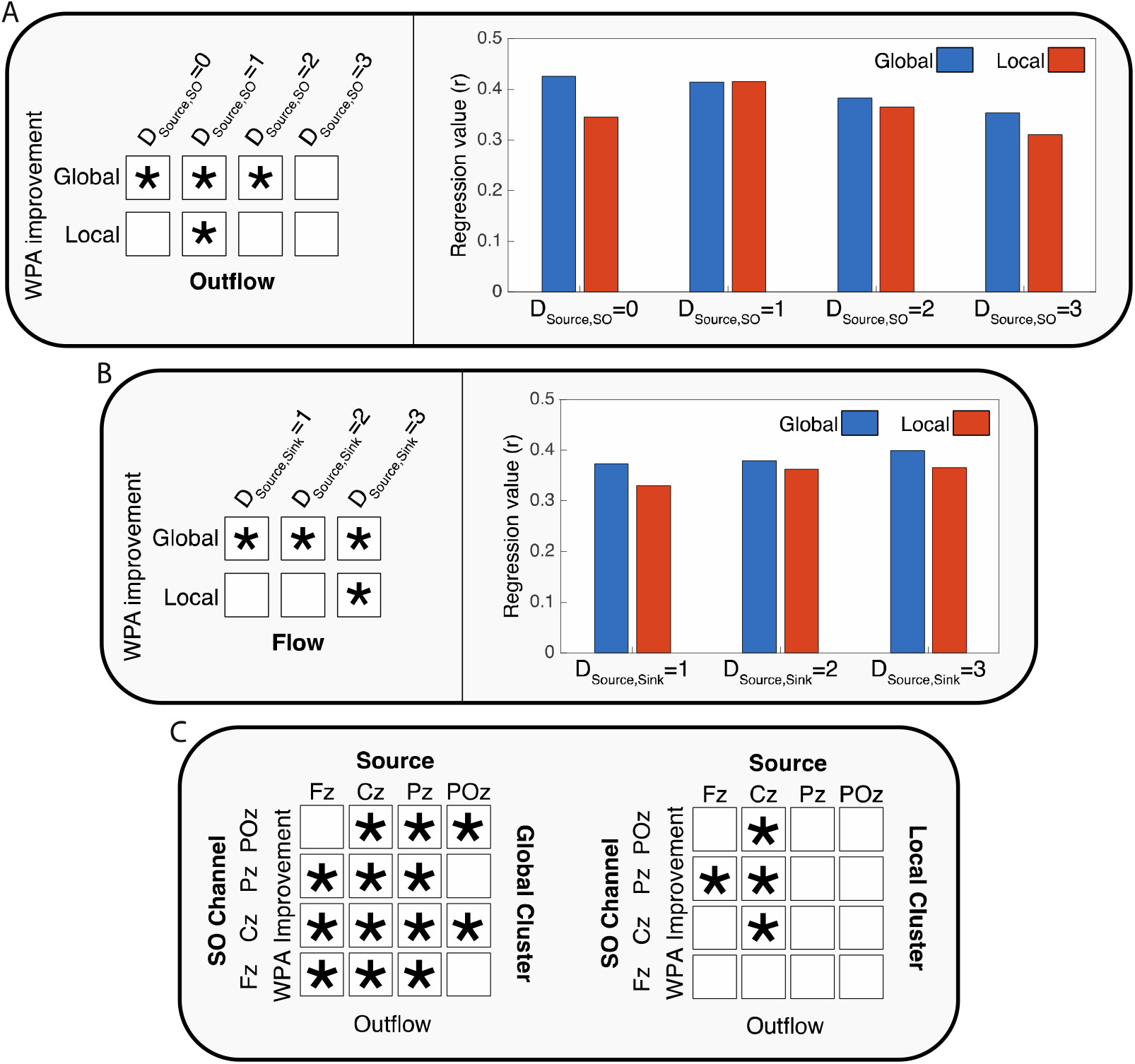
The effect SO clusters (local and global) on the relation between causal information flow and WPA improvement. The asterisk shows significant linear relationships. A. Results of correlation and regression tests for four distances between source and SO channels (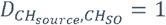 to 4). The left panel present the conditions that there was significant linear relation between outflow and WPA improvement (with p-values adjusted to Bonferroni correction). The right panel shows the r-values of regression in different conditions of distance of source and SO channel in the two clusters. B. Results of correlation and regression tests for three distances between sources and sinks 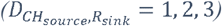 (with p-values adjusted to Bonferroni correction). C. Summarized results of the linear relations between outflow from source and SO channel combinations for global and local SO clusters (with p-values adjusted to false discovery rate (FDR)). Detailed results are presented in Figure S.9 and S.10.

Next, we averaged the outflows based on the distances between sources and SO channels in each cluster and examined correlations with WPA improvement. Given our prior result that short distances between source and SO channels have the highest information flow, we expected that global clusters with shorter distances between these channels would show the strongest associations between outflow and memory improvement. Indeed, our results showed significant relations between outflow in the global cluster when the distances between SO and sources are smaller than 4. In the local cluster, consistent with the reduced footprint of SOs in this cluster, we found a significant relation between outflow and WPA improvement only when the distance between SO and source was equal to 1 (Figure 5.A). Furthermore, r-value comparisons showed that the best condition (greatest r-value) for modeling WPA improvement was in the smallest distance between SO and source and for SOs in the global cluster.

We then conducted the same analysis between sink and source distances in each cluster. In each of the clusters, we calculated and averaged flows post-peak across distances between sink and source (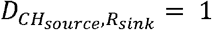 to 3). We found (Figure 5.B) a significant linear relation between flow and WPA improvement for all sink/source distances in the global cluster. Conversely, in the local cluster there was a significant relation between flow and WPA improvement only when sink and source were at the farthest distance 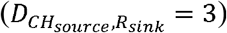. In sum, these results demonstrate that the best condition for modeling WPA improvement based on information flow is during global SOs across long-range neural channels.

## Discussion

In this work, we investigated temporal and topographical properties of the sleep slow oscillation (SO) by measuring effective connectivity during the SO phase, which establishes directional causality of information flow. We estimated information outflow from individual channels and flow between sources and sinks. First, consistent with prior findings, we found that information outflow at the SO trough and in non-SO windows are similarly dampened. Within this state of reduced communication, however, we identified two peaks of large-scale outflow before and after the SO trough. The direction and magnitude of the outflow peaks depended on the topographical position of the SO, dramatically decreasing with farther distances from the SO origin. Next, we considered source and sink quantifiers, and determined that sources closer to the origin of the SO were the biggest senders of information, whereas sinks farthest from the SO were the highest receivers. Cluster analysis confirmed that, compared to local SOs, global SOs mediated greater information flow. Taken together, our effective connectivity findings suggest that despite the generally disconnected state of NREM communication, SOs facilitate two bursts long-range causal communications across brain areas. As the goal of the study was to determine how the brain organizes network activity to support memory, we tested the temporal and topographic conditions of NREM sleep that predict overnight memory improvement. Our results show that memory improvement is predicted by conditions that maximize the distance between sink and source channels, relative to the SO origin, specifically during global SOs. Our findings are consistent with the notion that SOs provide the temporal framework for systems level communication which is a necessary condition for hippocampal, episodic memory consolidation.

### Generalized partial directed coherence as a Granger causality measure

Based on its mathematical definition, Granger causality can estimate the causal information flow between different time series, which is then interpreted as effective connectivity. The main aspect that differentiates effective connectivity from functional connectivity is its ability to quantify relative influences that are a-synchronous. Therefore, Granger causality can find “predictive causality” (Diebold, 2007) and identify the cause-effect relations with constant conjunctions (Maziarz, 2015). Within this concept, GPDC was used in this study to model the causal information flow between different regions that could be found in sleep EEG signals. GPDC as a generalized form of PDC can distinguish indirect and cascade connections from direct connections, based on partial coherence notation (Baccalá & Sameshima, 2001). PDC (and GPDC) will show propagation of information flow only when there is phase difference between sink and source, thus producing null information flow as result of factors that have no phase difference, such as volume conduction. In practice, volume conduction can have some minor influence on GPDC calculations, such as increasing the noise level. These influences are not critical (Blinowska, 2011), especially for our study, in which we have averaged the GPDC values over many samples (SOs).

With respect to the potential confound introduced by SOs being large amplitude objects, it is important to note that GPDC leverages weighted averaging to bypass amplitude difference between signals. Although greater amplitude in EEG signal can potentially represent enhanced synchrony in the neuronal activity giving rise to the signal, and hence can potentially be a source of information, the very definition of Granger causality between a sender-and-receiver of information pair involves a balance of the amplitude between signals, thus accounting for amplitude and rendering GPDC not sensitive to high amplitude effects.

### SOs shape the causal communication network in NREM sleep

Oscillatory activity has been linked to neural firing rate and spiking activity via multi- and single-unit recording and intracranial EEG (Nir et al., 2011). Although higher firing rate and spiking activity has been interpreted to signify greater potential outflow of information (Theunissen & Miller, 1995), the precise nature of the SO-triggered information flow has yet to be identified. Nir et al. (2011) reported that the mean firing rate of units within brain regions (e.g., orbitofrontal cortex, anterior cingulate, supplementary motor, parietal cortex, parahippocampal gyrus, and hippocampus) increased up to 200% 500ms before and after the SO trough, and decreased by 40% within the SO trough in comparison with the mean firing rate in NREM (N2 and SWS) (Nir et al., 2011). In the current work, we also found two peaks of information outflow (pre- and post-SO trough), however, the timing and phase of the outflow peaks in our study (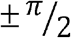 and about ±250ms from the SO trough) did not match the peak firing rate reported by Nir et al. (2011) (± π and about ±500ms from the SO trough), suggesting that there are other components and conditions which affect causal information flow in addition to spiking activity. While spiking activity can represent abundance of information in a source, availability of the sink to receive such information is a necessary condition to active communication, like causal information flow. Thus, despite a peak in neuronal firing rate 500ms before and after the SO trough, our data suggests that the best condition for information flow between sources and sinks is found closer to the SO trough, at about ±250ms delay.

Prior studies have demonstrated that brain communication is significantly modulated by the transition from wake to sleep. Using transcranial magnetic stimulation (TMS) to probe propagation patterns across wake and sleep stages, Massimini et al. showed the lowest levels of brain connectivity during NREM sleep (Massimini et al., 2005). Furthermore, effective connectivity analysis using functional magnetic resonance imaging (fMRI) and simultaneous EEG recordings have described N2 as an unstable network and SWS as a stable network (Kung et al., 2019). Together with our current findings of equivalently low levels of information outflow during SO troughs and non-SO windows during NREM, one emergent hypothesis is that causal communication across brain regions during NREM sleep is generally poor, but punctuated by bursts of selective high-communication events, mediated by SOs.

### Causal information outflow depends on distance from SO channel

Our results demonstrate several key properties of the SOs that contribute to their communication profile. Along with time and phase of the SO gating causal information outflow, our results demonstrate that SOs enable information flow with topographic specificity as well. In particular, the distance between source and SO channels had the strongest effect on peak height (i.e., amount of information flow). Thus, the closer the source, or information sender, is to the SO channel, the higher the information outflow. These peaks were not identical before and after the SO trough, suggesting that brain communication differs during the down-to-up transition compared with the up-to-down transition of the SO. One potential explanation for these differences may be that we combined SOs of different types, including single and sequential SOs (Mölle et al., 2011), wherein sequential SOs have pre- and post-SO peaks that are temporally linked with each other. In these cases, the pre-peaks may be influenced by the preceding SO and the current SO, whereas post-trough peaks would be only affected by the current SO. In agreement with our findings, studies have shown functional differences between pre- and post-SO activity, including greater synchronization between SOs and spindles (Hahn et al., 2020; Niknazar et al., 2015) and more efficient cueing during targeted memory reactivation (TMR) during the post-SO trough peak (Göldi et al., 2019). Thus, our data support the notion that there are functional and effective connectivity differences between SO peaks, but that further research is needed to tease apart properties of single or sequential SOs. Lastly, by modeling the outflow peaks, we showed a significant effect of the SO channel, with the positive coefficient demonstrating an increase in the height of outflow from anterior to posterior regions. These results are likely due to the presence of significantly fewer SO in posterior regions, suggesting the intriguing possibility that total information flow is not higher in the posterior channel in general, but rather that during the few posterior SOs that do occur, there is a higher amount of information outflow than other areas. Future research may determine why outflow increases as the number of detected SOs diminishes.

### SOs facilitate causal communication between more distant regions

The SO has also been described as a global traveling wave that recruits the entire cortical network (Neske, 2016). We, therefore, probed information flow by considering source and sink channels independently and together as quantifiers and discovered that regions farther from the SO channel are more engaged in receiving information. This suggests that global brain communication during the SO may be organized with local areas near the SO sending more information and local areas farther from the SO receiving more information. As such, brain areas engaged in high causal information outflow (i.e., sending information) have a low chance of simultaneously receiving information. Given these results, we hypothesized that the effect of distance between sinks and sources and greater causal communication between farther regions should be specifically accentuated in the case of global SOs which involve a greater number of scalp locations (Malerba et al., 2019). We tested this hypothesis by using global or local SOs as predictors in our model and demonstrated a significant effect of SO cluster on the height of the flow peaks, with global SOs having higher peak heights than local SOs.

### SOs as a facilitator for long-term memory

One leading hypothesis of memory consolidation suggests that encoding networks get reactivated during sleep (Foster, 2017), specifically during NREM sleep (Diekelmann & Born, 2010b). Given the near-infinite diversity of the encoding networks, memory reactivation requires both short and long-range information transfer across a wide range of brain areas. It is well established that SOs provide temporal coordination for memory-related brain activity during systems consolidation (Diekelmann & Born, 2010b) and a greater effect of TMR has been shown when cues are coupled with the down-to-up phase of the SO (Göldi et al., 2019; Ngo et al., 2013), similar to the post-peak in the current results. However NREM is a sleep period generally characterized as showing a breakdown in connectivity with increasing depth (Massimini et al., 2005), which would not be conducive to reactivation.

Our results pose a potential solution to this seeming paradox, with global SOs providing the temporal-topographical event framework whereby long-range information flow increases dramatically compared to the relatively local activity patterns of the rest of the sleep period. Specifically, as memories get reactivated in cortical-hippocampal networks, global, traveling SOs mediate systems level communication across all source-sink pairs, even for sources far from the SO origin and close sink and source pairs. In agreement with this finding, for local SOs, the relation between causal flow and episodic improvement was limited to sources closest to the SO channel and long-distance pairs of sinks and sources.

By describing the space-time dynamics of sleep oscillations, we can non-invasively acquire a window on the coordination and brain processing patterns of basic psychological function as well as in cognitive disorders or conditions that currently lack a mechanistic connection between their understood origins and their cognitive manifestations. Intriguingly, we have shown that SOs coupled with sleep spindles increase causal information flow. Future investigations should test how nested oscillations (e.g., SO-spindle complexes) promote greater information flow and their functional relevance for memory consolidation.

### Limitations and suggestions

Limitations of this study include combining all SOs in the calculation of causal information flow ignoring distinctions between single or sequential SOs, which may have affected the results in pre-vs post-peaks. Our Granger causality analysis was applied to sensor level data (EEG signals), which is clearly not optimal when talking about directed connectivity among neuronal sources. In principle, one could use a source reconstruction technique to create a set of virtual electrodes and then repeat the Granger causality analysis in source space. This would lend greater validity to our conclusions, in relation to the underlying cortical connectivity. Also, we have referred to GPDC (a Granger causality analyses) as effective connectivity to emphasize its causal nature. However, strictly speaking, Granger causality is a directed form of functional connectivity, because it rests upon statistical associations. A more complete analysis would be dynamic causal modelling of complex cross spectra; effectively parameterizing a neural network in source space and then finding the effective connectivity that best explains the Granger causality. The advantage of this kind of modelling is that it can resolve recurrent effective connectivity. In addition, we focused on frontal to occipital brain regions ignoring temporal regions, which may affect interpretation of results. For each region, we considered three EEG channels that contributed to region’s estimate. It is possible that increasing the number of channels included in each region (i.e., increasing scalp resolution) may produce more robust results. Also, considering a larger number of regions (e.g., Fronto-Central, Centro-Parietal, and Parieto-Occipital) may improve the efficiency of modeling the relation between areas. In this study, the simplest approach for attributing values to the distance of channels and regions was adopted (discrete linear values). The effect of these distances in our estimates could alternatively be modeled with real-world values of the distances across the brain cortex. GPDC as a Granger causality estimator -consistent with other methods in this family-represents effective connectivity in the frequency domain. In an effort to reduce complexity, we chose to average the effective connectivity values, supported by the lack of mechanistic reasons to preferentially concentrate on a specific a-priori frequency range for directional connectivity estimates. Future studies should address potential differences in effective connectivity profiles when GPDC is evaluated only in specific frequency bands, possibly concentrating on ranges that are known for their physiological relevance and interaction with SOs, such as sigma.

