## supplementary for "Slow oscillations promote long range effective communication: the key for memory consolidation in a broken down network"

**Materials and Methods**

**Data**

Overnight sleep was measured in 59 healthy adults (20.5 ± 2.57 years, 26 females) with no history of psychological and neurological problems. All participants signed informed consent, which was approved by the University of California, Riverside Human Research Review Board. EEG signals were acquired using 32-channel cap (EASEYCAP GmbH) with Ag/AgCI electrodes placed according to the international 10-20 System at sampling rate of 1000 Hz. 22 EEG channels were used for EEG signal recording and the others were used for reference, ground and other biosignals including EMG, EOG and ECG. EEG signals were re-referenced to the contralateral mastoid (A1 and A2) and down-sampled to 256 Hz after the recording. Raw data were visually scored in 30-sec epochs into Wake, Stage 1, Stage 2, Slow Wave Sleep (SWS) and rapid eye movement (REM) sleep according to the Rechtschaffen & Kales’ manual (Kales et al., 1968) using HUME (Saletin & Greer, 2015), a custom MATLAB toolbox. All the 22 EEG channels were used for SO detection and the clustering process. Effective connectivity was computed considering 12 EEG channels (F3, Fz, F4, C3, Cz, C4, P3, Pz, P4, O1, POz and O2) which were filtered with a Butterworth high pass filter with stop band of 0.5 Hz and order of 46.

To investigate the relation between causal information flow and improvement in long-term memory (LTM) the word pair association (WPA) test was deployed. The subjects were divided into two groups randomly. The first group completed the WPA encoding session before the night of sleep and the second group completed the WPA encoding session in the morning followed by night of sleep. As we were interested in causal information flow in SOs and its effect on improvement of LTM, the performance in the pre-sleep (test 1 in the first group and test 2 in the second group) and post-sleep (test 2 in the first group and test 3 in the second group) test sessions of WPA task were used and the ratio of post-sleep performance to pre-sleep performance was considered as sleep related WPA improvement ratio (Figure S.1).

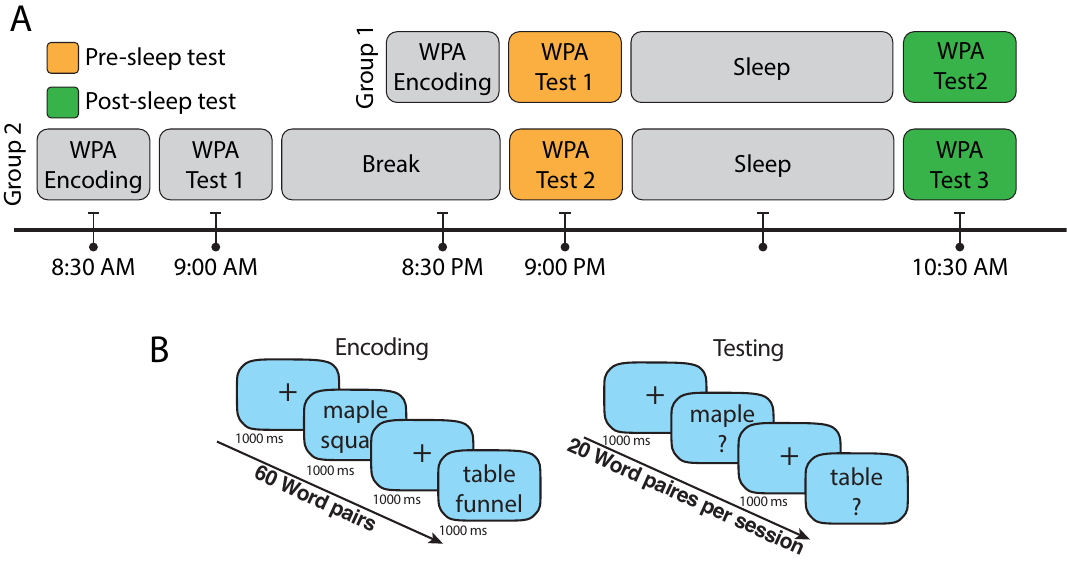

Figure S. 1 Timeline of the encoding and test sessions of WPA task and details of the encoding and test sessions. A. Participants in group 1 (n=26 after excluding participants who did not complete all the task sessions) reported to the laboratory in the evening and completed the encoding session of WPA and immediate testing (Test 1, pre-sleep) at around 9 PM and prepared for sleep. They were woken up at the morning and after a breakfast completed the next testing session (Test 2, post-sleep) at 10:30 AM. In group 2, participants (n=26 after excluding participants who did not complete all the task sessions) completed the WPA encoding and immediate testing session (Test 1) at around 8:30 AM. Then, participants left the lab to continue their day and came back to the lab to complete the second testing session (Test 2, pre-sleep) at around 9 PM. In the next day morning, participants were woken up at 9 AM and after a breakfast completed the last test (Test 3, post-sleep) at 10:30 AM. B. In the encoding session of the WPA task, 60 pairs of words with length of 3-9 letters were presented vertically stacked and shown twice in random order. Each word pair was presented for 1000ms and followed by a fixation cross for 1000ms. Immediately after encoding, subjects were trained to criterion using a test in which participants were shown one word of the pair and were required to type in the associated word. Incorrect trials were repeated and feedback was provided. The participants had to achieve 70% accuracy to finish the encoding session. The used word pairs in the encoding session were divided into 3 groups of 20 word pairs for the testing sessions. In each testing session, participants were shown one word of the pair and were required to type the associated word.

**SO detection**

We used an SO detection algorithm introduced by Dang-Vu et al. (Dang-Vu et al., 2008) to find SOs in stage 2 and SWS, measuring zero-crossing points of filtered EEG signal in the 0.1-4Hz range. Four criteria were used to identify an SO: 1) The wave minimum was below or equal to 80 uV; 2) The range of values between maximum and minimum voltage was at least 80 uV; 3) The time between the first and second zero crossing in the data had to be between 300ms and 1s; 4) The total duration of the candidate event was at most 10s. The time margin of detected SOs from the Fz, Cz, Pz and POz channels were used for effective connectivity computation, all detected SOs in all 22 EEG channels were used for the clustering process.

**Effective connectivity estimation**

Granger causality is an analytical definition of causality between two time series. Intuitively, according to this definition, time series $x_{1}$ Granger-causes $x_{2}$ if using past values of $x_{1}$ ($t<t_{0}$) improves The prediction of the current value of $x_{2}$ ($t=t_{0}$) (Granger, 1969). This principle can be modeled via a multivariate autoregressive (MVAR) model. MVAR models the interactions between past values of $N$ signals ($X$) in relation to the present time values ($X\left( n \right)=\left\{ x_{1}\left( n \right), x_{2}\left( n \right), \ldots, x_{N}\left( n \right) \right\}$) using a linear regressive model (Eq.1).

(1)

$$X\left( n \right)=\sum_{k=1}^{p} A_{k}X\left( n-k \right)+w\left( n \right)$$

Where $X\left( n \right)$ is a vector containing amplitudes of all signals in time, $n$ marks the time stamp at which $X$ is being computed, and $p$ is the model order (i.e., how many steps back in time are used to model the current values). $A_{k}$ is the matrix of MVAR coefficients and each element $a_{ij}\left( k \right)$ represents how much the *j-*th signal at time $n-k$ affects the $i$th signal at time $n$*.* $w\left( n \right)$ is the model’s additive Gaussian noise vector, with zero mean and covariance matrix $\Sigma$*.* Several measures use MVAR to estimate Granger-causality-based effective connectivity, such as the Granger causality index (GCI), the directed transfer function (DTF), partial directed coherence (PDC) and numerous variations of these measures (Blinowska & Kamiński, 2006). PDC measures the directed linear relation between two time series using MVAR coefficients in the frequency domain (Eq.2).

${PDC}_{ij}\left( f \right)=\frac{A_{ij}\left( f \right)}{\sqrt{\sum_{k=1}^{N} \left| A_{kj}\left( f \right) \right|^{2}}}$

(2)

Where $f$ is normalized frequency [0 to 0.5] Hz (here based on the sampling rate of 256 relative to actual frequency range of 0 to 128 Hz) where $A_{ij}\left( f \right)$ is the MVAR coefficients in frequency domain (Eq.3). In this study, the frequency is sampled at 1/128 Hz relevant to 1 Hz in actual frequency.

$A_{ij}\left( f \right)=\delta_{ij}-\sum_{k=1}^{p} a_{ij}\left( k \right)e^{-2j^{*}\pi fk}$

(3)

(3)

For $\delta_{ij}=1$ whenever $i=j$ and $\delta_{ij}=0$ otherwise, and $j^{*}=\sqrt{-1}$.

PDC can be affected by unbalanced signals amplitude. Therefore, its generalized version (GPDC) introduces a re-definition that improves PDC’s estimations in scenarios with severely unbalanced errors caused by unbalanced amplitude scales (Baccala et al., 2007). Specifically, GPDC uses elements of noise covariance matrix ($\Sigma$) to overcome this potential bias(Eq.4).

$\bar{\pi}_{ij}\left( f \right)=\frac{\frac{1}{\Sigma_{ii}}A_{ij}\left( f \right)}{\sqrt{\sum_{k=1}^{N} \frac{1}{{\Sigma_{kk}}^{2}}\left| A_{kj}\left( f \right) \right|^{2}}}$

(4)

Where $\bar{\pi}_{ij}\left( f \right)$ is the matrix of causal information flow (GPDC), in which the j-th column estimates causal information outflow from the j-th signal to all other signals, while $N$ counts the total number of signals considered. The values are normalized over each column separately therefore, outflows in each column are comparable only within the specific column.

For each considered SO event in each of Fz, Cz, Pz and POz channels, the time interval of 1s before to 1s after the negative trough of the SO was considered for the analysis. To compute causal information flow during SOs, GPDC were computed in sliding windows for each SO separately, which allowed us to follow causal information flow changes over time. The window length (w) was 0.5s to maintain a relative stationarity of EEG signals and stride was 7.8ms. Therefore, we estimated 193 GPDC three-dimensional tensors over the time course of each SO. Akaike information (Aho et al., 2014) was used to choose the appropriate model order $p$, resulting in $p=13$ selected for all the windows. To reduce the complexity of computations and having more robust results, the GPDC tensors sized $N\times N\times N_{Frequency}$ were averaged over the frequencies, to construct the GPDC matrix $\pi\left( i,j \right)$. Also, to reduce variance, causal information outflow from a given channel (Fz, Cz, Pz and POz) to all channels in each other areas including frontal, central, and parietal were averaged. Figure 1.A represents the process of estimating the outflow to each area. 12 channels of EEG signals ($N$=12) were considered for computing GPDC matrices (F3, Fz, F4, C3, Cz, C4, P3, Pz, P4, O1, POz and O2). Fz, Cz, Pz and POz were considered sources of causal information in estimating effective connectivity, and outflow from these selected central channels to all 12 channels was quantified for causal information flow. We defined two types of outflow quantifiers, one that combined the outflow to all channels in regions different than the one that included the source, and one that concentrated on a given sink region. Their respective equations are:

(5)

$${CH}_{Outflow}= \sum_{i\in D} \left( \pi\left( i,CH \right) \right)^{2}, D=\left\{ all channels excluding channels in the same region as CH \right\}$$

$CH\to R= \sum_{i\in R} \left( \pi\left( i,CH \right) \right)^{2}, R=\left\{ all channels in the R region \right\}$.

Here, $CH$ is one of the four source channel and $R$ is one of the sink regions (frontal (F) including [Fz, F3 and F4] channels, central (C) including [Cz, C3 and C4] channels, parietal (P) including [Pz, p3 and P4] channels and occipital (O) including [POz, O1 and O2] channels). These quantifiers were computed for each window of SOs and averaged in different conditions, including SOs in each source channel separately and different pairs of source channel and sink region.

$\pi\left( i,j \right)$ matrices were calculated for each SO in Fz, Cz, Pz and POz channels as the source of causal information flow channels. By using the sliding window technique, for each 500ms window, the quantifiers (${CH}_{Outflow}$ and $CH\to R$) for all four source channels and sink regions were calculated (Figure 1.A). The resulting values were assigned to the central time of their respective window. Therefore, for each SO in a source channel and each quantifier there were 193 values between 750ms before to 750ms after the SO trough. For each participant, each quantifier was averaged over all SOs in each of Fz, Cz, Pz and POz channels separately, producing a time series per each quantifier per channel.

To access the quantifiers variation over the SO’s phase we used nonlinear resampling in three steps. In the first step, the phase of each SO was estimated using the Hilbert transform between 1s before and 1s after the SO trough (whose phase was assigned as zero as it is represented in Figure 1-B). In the next step, we selected a valid time range by including only the times in which the phase increased monotonically between -pi and pi. In the last step, we generated a phase series in the valid time range increasing from -pi to pi with fixed step at pi/64 by re-sampling the quantifier within the valid times. Although the proposed method starts with the SO trough as the timing reference, this referencing is immediately lost after resampling from the time to the phase domain. Thus, any exact timing reference is removed in favor of matching all pre- and post-trough peaks to their phases. For example, the computed quantifiers of the windows centered at positive peak of EEG before the SO troughs in Fz channel have been averaged such that their phases are the same (-π).

**Clustering**

The same approach as (Malerba et al., 2019) was used to cluster SOs into global and non-global clusters. Briefly, for each detected SO in each of the 22 channels, a binary vector $V$ with the length of the number of all channels was constructed (Eq.6).

(6)

$$V=\left[ c_{1}, c_{1},\ldots, c_{22} \right]$$

Where $c_{j}$ was assigned 1 if there was an SO in j-th channel in the time range of ±400ms of SO’s trough time in the i-th channel and 0 if there was not an SO in that time margin in j-th channel. Next, a K-means algorithm with Hamming distance was run to categorize all vectors into two clusters (K=2). The K-means algorithm was run with 200 repetitions minimizing within-cluster distance, in both stage Stage 2 and SWS.

**Statistical Analysis**

A linear mixed-effects (LME) model was used to model the GPDC causal information flow peaks and investigate the effect of different variables, including source channel, sink region, etc. The LME is an extension of linear regression models for data that are collected and summarized in groups (Jiang, 2007). LME models describe the relation between a response variable and the independent variables, with coefficients that can vary with respect to one or more grouping variables. Each participant was considered as a random effect and each variable was considered as an independent fixed effect. After fitting an LME model to the data, multiple values were considered, including the coefficients of the model and p-value related to each parameter. The p-value of each fixed effect was interpreted as a significant linear effect of the variable on the model. To investigate the predictors as independent fixed effects we assigned numbers to predictor values using a linear approach. Number one to four were used for the anterior channel or region (Fz or frontal region) to the posterior channel or region (POz or occipital region). The same numbers were used to represent the distance between channels and regions. Also, to model the effect of the phase on the amplitude of the peaks, the pre-peak of the flow was identified with the value one and the post-peak of the flow was marked with two.

Analysis of variance (ANOVA) was used to compare quantifier values in each peak and the non-SO windows. Also, a one-way ANOVA was performed to test significance between different time windows during SOs and non-SO, and post-hoc analysis was applied to identify windows that were significantly different from the non-SO windows.

Linear model was used for correlation and regression test to investigate the linear relation between WPA improvement ratio and causal information flow in different types of conditions. Three types of condition were considered to group and average the causal information flow in each condition and fit the linear model using the averaged flow as the predictor and WPA improvement ratio as the dependent variable. In the first type of condition, the distance between SO and source channel ($D_{\mathrm{CH}_{\mathrm{source}},\mathrm{CH}_{\mathrm{SO}}}$) was considered as the condition in which the distance could be 0 to 3 (total 4 condition groups). The distance between source channel and sink region ($D_{{CH}_{source},R_{sink}}$) was considered as the second condition type with value of 1 to 3 (total 3 condition groups). The third condition type was considered as relative distance between sink and source with respect to SO channel which was defined as $D_{R_{\mathrm{sink}},\mathrm{CH}_{\mathrm{SO}}}-D_{\mathrm{CH}_{\mathrm{source}},\mathrm{CH}_{\mathrm{SO}}}$ and condition groups were defined as positive, negative and zero values (total 3 condition groups). The positive values represent the condition where the source is closer to SO channel than the sink to SO channel and the negative value shows the opposite condition. In all the statistical tests, mean±2std were considered as the outliner boundary and data outside of this margin were excluded.

**Supplementary figures and tables**

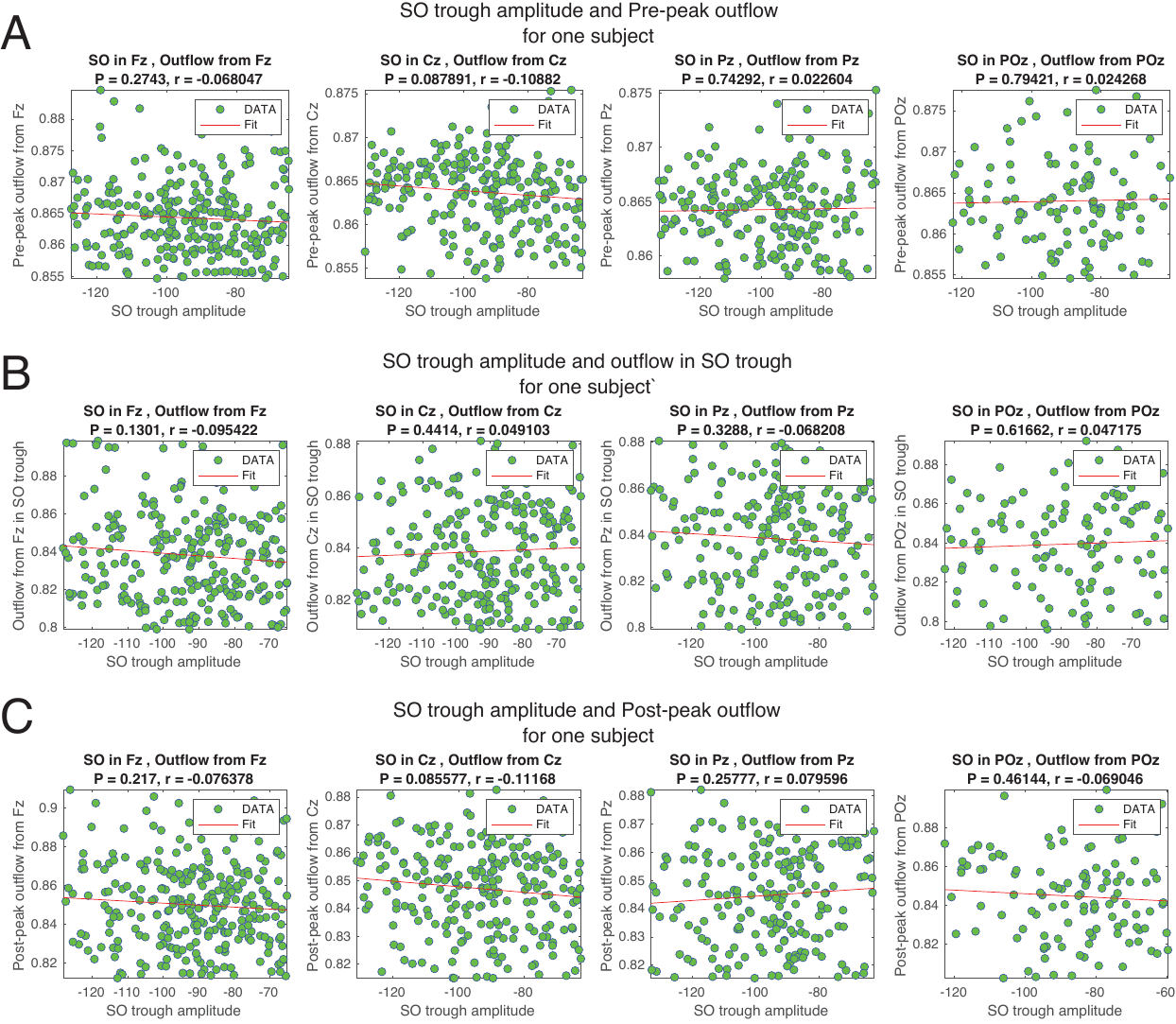

Figure S. 2 Correlations between SO amplitudes and information outflow using all SOs of an example subject in each of the channels. A) Correlations between SO trough amplitudes and pre-peak (phase = -π/2) outflow. B) Correlations between SO trough amplitudes and outflow in SO troughs (phase = 0). C) Correlations between SO trough amplitudes and post-peak (phase = π/2) outflows. No significant relations between SO amplitudes and outflows in different phases were found.

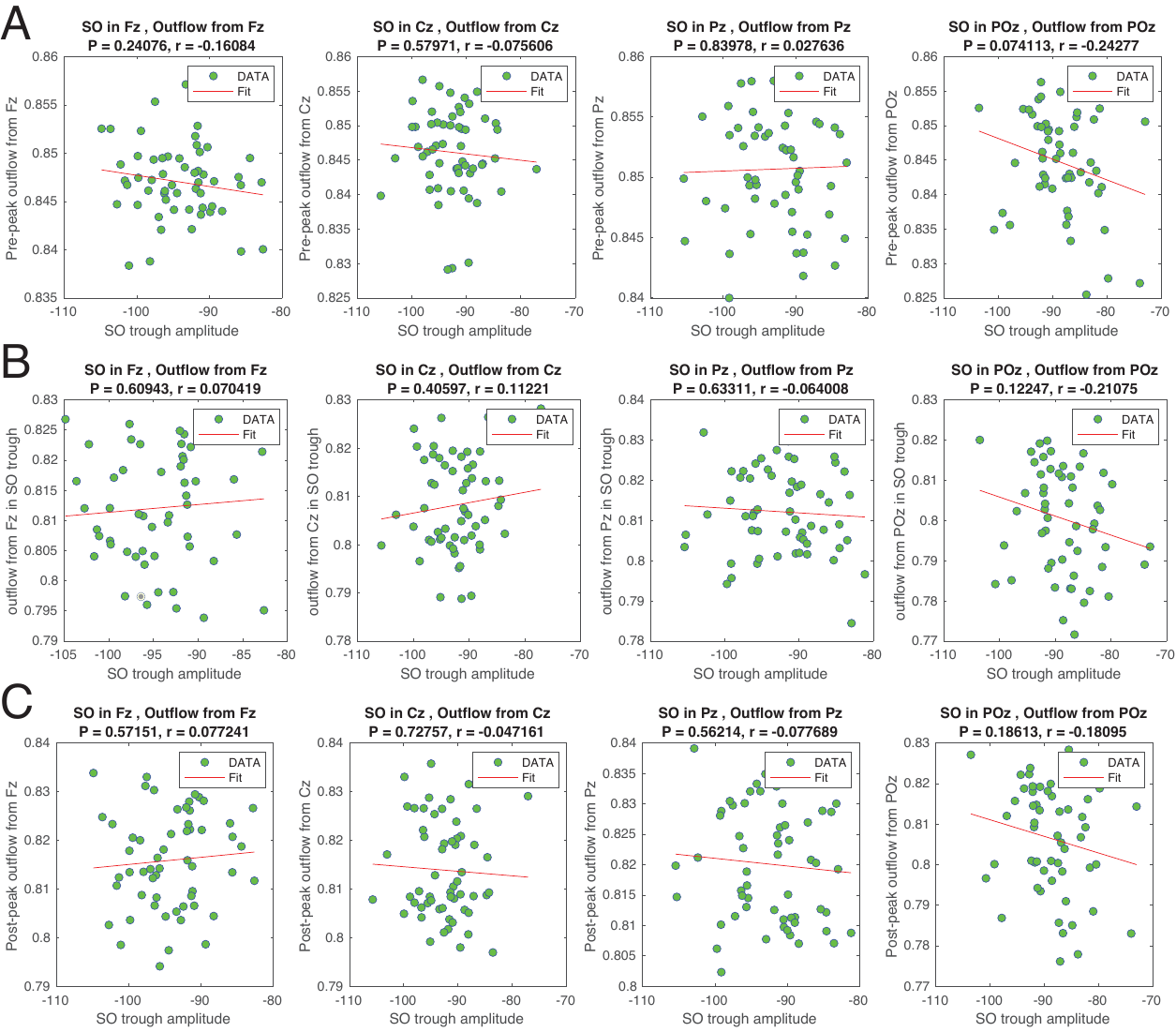

Figure S. 3 Correlations between SO amplitude averages for each subject and information outflows in each of the source/SO channels separately. A) Correlations between SO amplitudes and pre-peak (phase = -π/2) outflows. B) Correlations between SO amplitudes and outflows in SO trough (phase = 0). C) Correlations between SO amplitudes and post-peak (phase = π/2) outflows. No significant relations were found between SO amplitudes and outflows in different phases considering all the subjects.

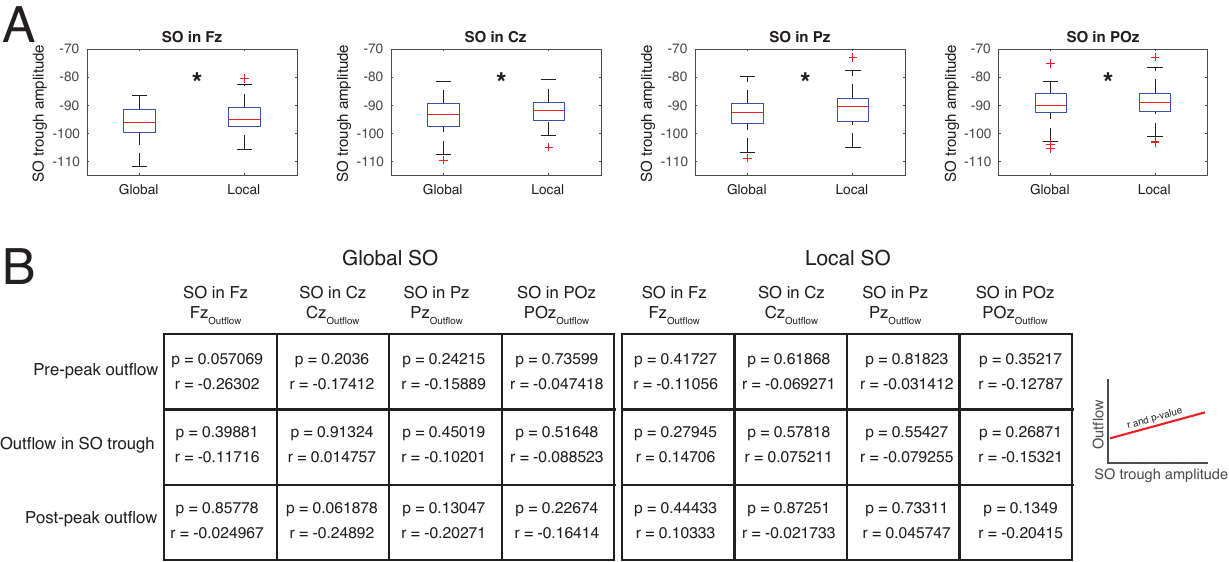

Figure S. 4 A) Distribution of SO amplitudes in each of the channels and in each of the two clusters (Local and Global). SOs in the Global clusters had greater amplitude (absolute value) than the Local cluster. B) The same analyses as Figure S.3 for SOs in each of the clusters separately. The results showed no significant relation between SO amplitudes and outflows in either cluster.

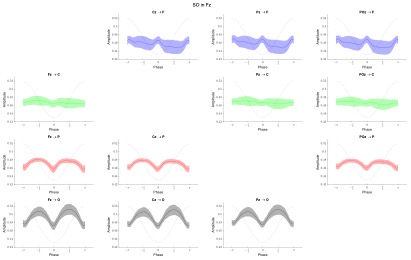

Figure S. 5 $CH\to R$ quantifiers for each of the source channel–sink region when SO channel is Fz. Each row represents the flow phase series to specific sink region. Each column represents flow phase series from each of the source channels.

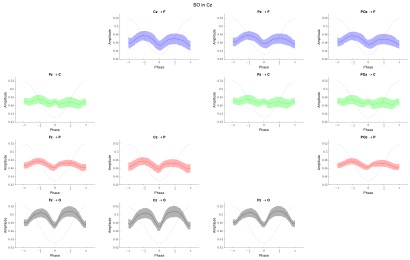

Figure S. 6 $CH\to R$ quantifiers for each of the source channel–sink region when SO channel is Cz. Each row represents the flow phase series to specific sink region. Each column represents flow phase series from each of the source channels.

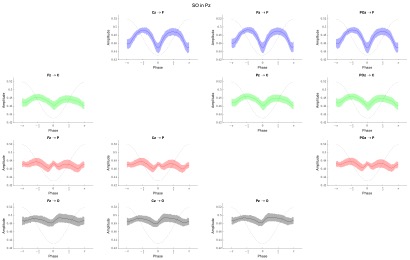

Figure S. 7 $CH\to R$ quantifiers for each of the source channel–sink region when SO channel is Pz. Each row represents the flow phase series to specific sink region. Each column represents flow phase series from each of the source channels.

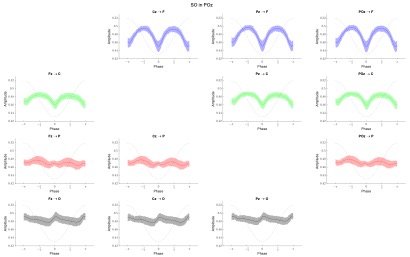

Figure S. 8 $CH\to R$ quantifiers for each of the source channel–sink region when SO channel is POz. Each row represents the flow phase series to specific sink region. Each column represents flow phase series from each of the source channels.

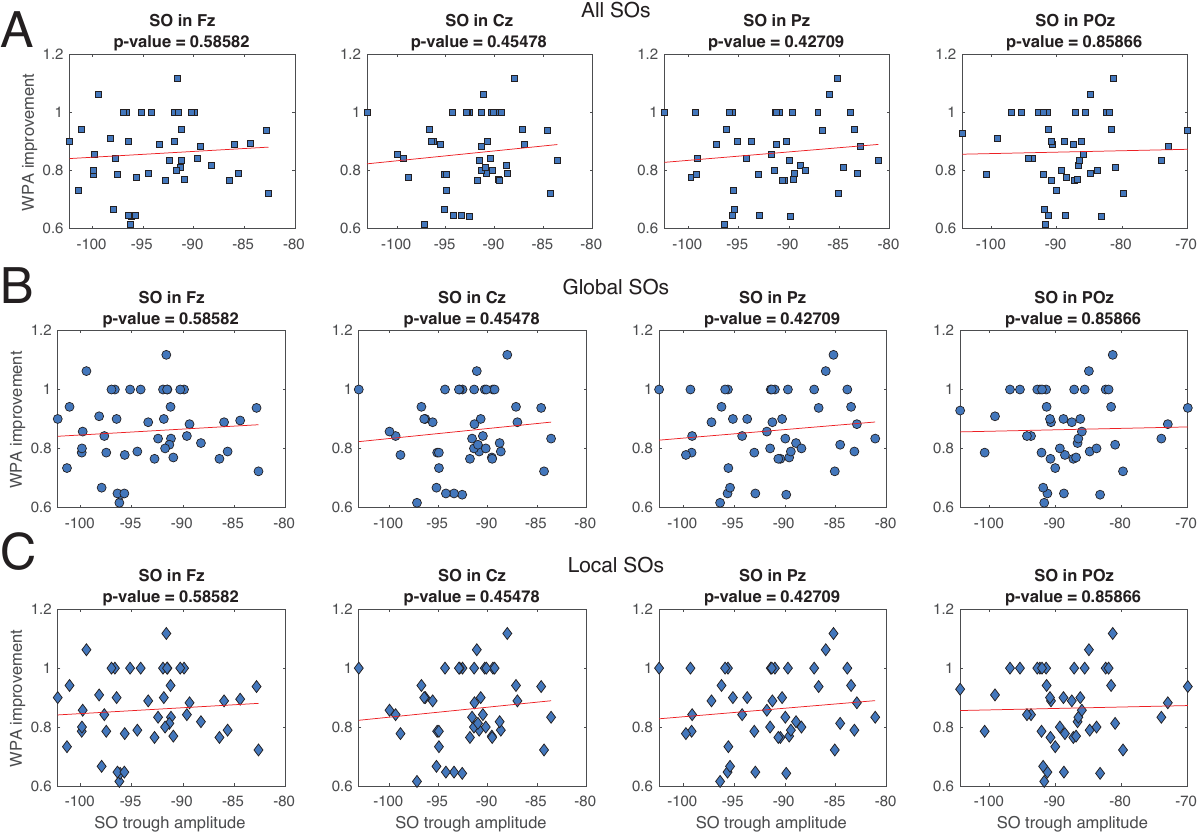

*Figure S. 9* *Correlations between SO amplitude and WPA improvement in each of the source/SO channels separately. A) Correlations between all SO amplitudes and WPA improvement (no significant effects). B) Correlations between Global SO amplitude and WPA improvement (no significant effects). C) Correlations between Local SO amplitude and WPA improvement (no significant effects).*

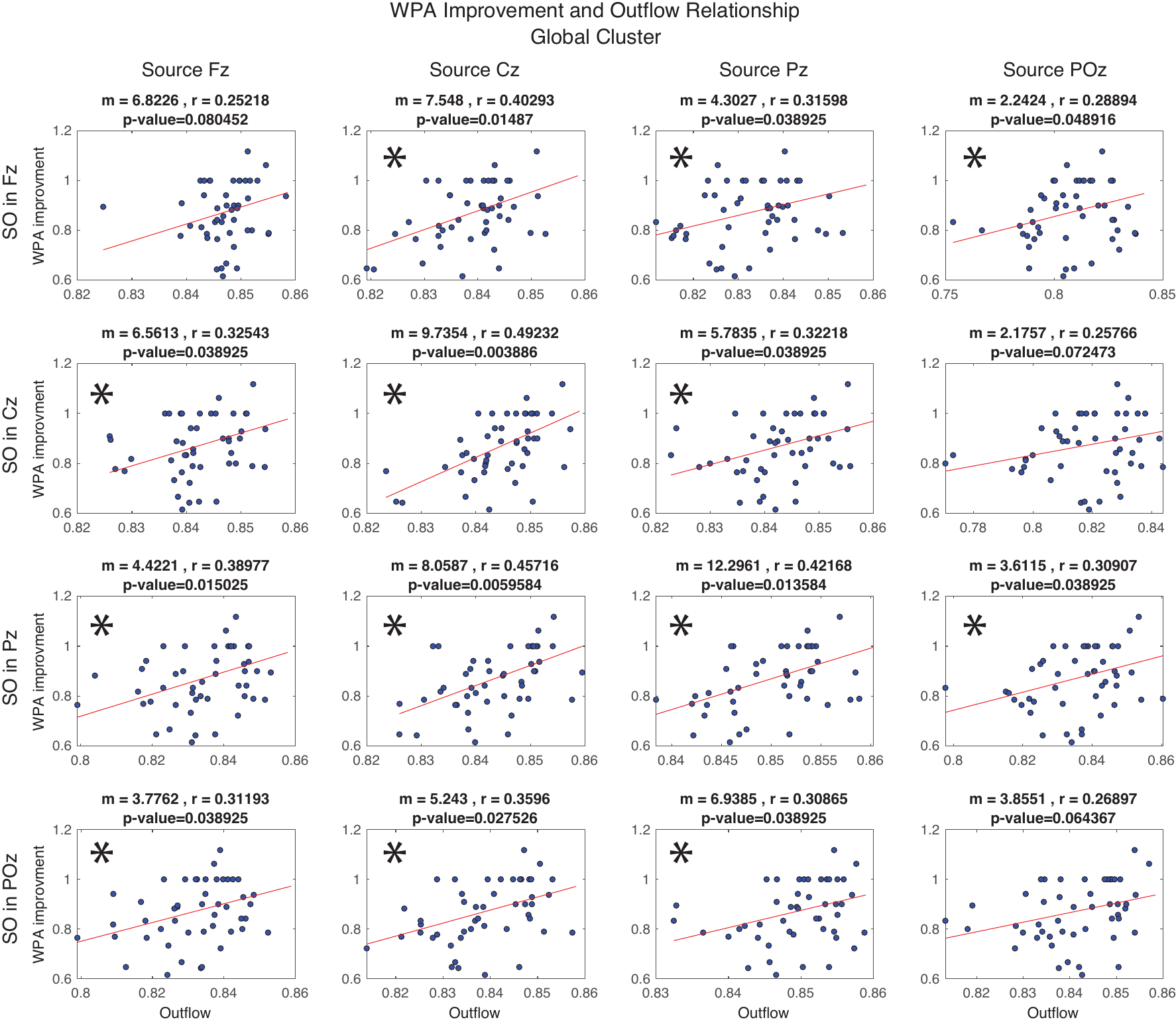

Figure S. 10 ) The results of the testing linear relationship between outflow from each source channel in different conditions of SO channel for the global SO cluster. Each row and column represents condition of SO and source channel respectively. The significant linear relationships are marked with asterisk (with p-values adjusted to FDR).

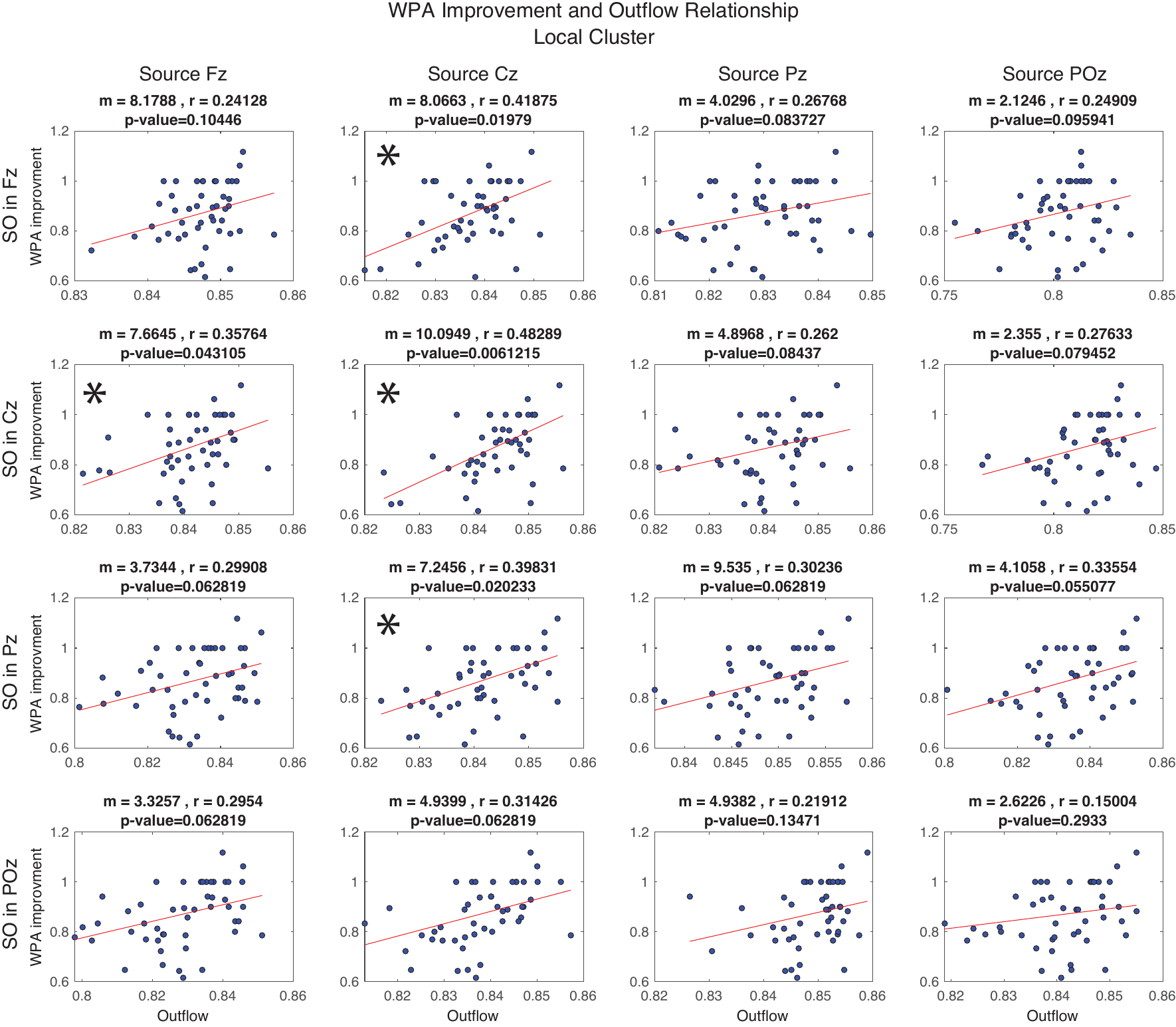

Figure S. 11 The results of the testing linear relationship between outflow from each source channel in different conditions of SO channel for the local SO cluster. Each row and column represents condition of SO and source channel respectively. The significant linear relationships are marked with asterisk (with p-values adjusted to FDR).

Table S. 1 Coefficients and p-values for linear predictors in LME models for amplitude of the outflow peaks.

|  | **Outflow peaks** | |
| --- | --- | --- |
| **Linear predictor** | Coefficient | p-value |
| SO channel | 0.0022 | **<0.0001** |
| Source channel | -0.0019 | 0.0532 |
| $D_{{CH}_{source},{CH}_{SO}}$ | -0.0054 | **<0.0001** |
| Peak phase (pre-/post-SO trough) | -0.0038 | **<0.0001** |

Table S. 2 Coefficients and p-values for linear predictors in LME models for peak information flow amount in SO phases $\pm\frac{\pi}{2}$. $D_{{CH}_{source},{CH}_{SO}}$ is distance between source channel and SO channel, $D_{R_{sink},{CH}_{SO}}$ is distance between sink region and SO channel, and $D_{{CH}_{source},{CH}_{R_{sink}}}$ is distance between source channel and sink region.

|  | **Flow peaks** | |
| --- | --- | --- |
| **Linear predictor** | Coefficient | p-value |
| SO channel | 0.0017 | 0.1457 |
| Source channel | 0.0002 | 0.2211 |
| Sink region | 0.0038 | 0.0543 |
| $D_{{CH}_{source},{CH}_{SO}}$ | -0.0006 | **<0.0001** |
| $D_{R_{sink},{CH}_{SO}}$ | 0.0076 | **<0.0001** |
| $D_{{CH}_{source},{CH}_{R_{sink}}}$ | 3.86e-05 | 0.8292 |
| pre-/post-peaks | -0.0029 | **<0.0001** |

Table S. 3 Coefficients and p-values for linear predictors in LME models for amplitude of the flow peaks after adding the cluster as a fixed effect.

|  | **Flow peaks** | |
| --- | --- | --- |
| **Linear predictor** | Coefficient | p-value |
| SO channel | 0.0015 | 0.1421 |
| Source channel | 0.0001 | 0.2910 |
| Sink region | 0.0033 | 0.077 |
| $D_{{CH}_{source},{CH}_{SO}}$ | -0.0005 | **<0.0001** |
| $D_{R_{sink},{CH}_{SO}}$ | 0.0067 | **<0.0001** |
| $D_{{CH}_{source},{CH}_{R_{sink}}}$ | 1.65e-05 | 0.9042 |
| Cluster | -0.0008 | **<0.0001** |
| peak phase (pre- or post- SO trough) | -0.0026 | **<0.0001** |

Table S. 4 Coefficients and p-values for linear predictors in LME models for amplitude of the flow peaks after adding SO-spindle coupling as a fixed effect.

|  | **Flow peaks** | |
| --- | --- | --- |
| **Linear predictor** | Coefficient | p-value |
| SO channel | 0.0016 | 0. 0854 |
| Source channel | 0.0001 | 0. 2364 |
| Sink region | 0.0035 | 0.0686 |
| $D_{{CH}_{source},{CH}_{SO}}$ | -0.0005 | **<0.0001** |
| $D_{R_{sink},{CH}_{SO}}$ | 0.0065 | **<0.0001** |
| $D_{{CH}_{source},{CH}_{R_{sink}}}$ | 1.65e-05 | 0.7205 |
| Cluster | -0.0008 | **<0.0001** |
| peak phase (pre- or post- SO trough) | -0.0027 | **<0.0001** |
| SO-spindle coupled | 0.0008 | **<0.0001** |
